# Small-scale habitat components as key drivers of biodiversity in urban park design

**DOI:** 10.64898/2026.06.30.735471

**Authors:** Gema Trigos-Peral, Joaquín L. Reyes-López

**Affiliations:** Museum and Institute of Zoology, Polish Academy of Sciences, Twarda 51/55, Warsaw, 00-818, Poland; Ecology Area, University of Cordoba, Edificio C-4 “Celestino Mutis”, Campus de Rabanales, Ctra, Madrid, Km. 396, 14071 Cordoba, Spain

**Keywords:** ant diversity, management, global change, urbanization, ecological refuge, species turnover, invasions, Argentine ant

## Abstract

Urban green spaces are increasingly recognised as important refuges for biodiversity, yet their ecological value depends strongly on design and management. Here, we investigate how fine scale structural and microhabitat components shape urban ant assemblages, using ants as indicators of broader arthropod responses to urbanisation. Ant communities were sampled in twelve urban green spaces in Córdoba (southern Spain) over a ten□year period (2004–2013) using pitfall traps, alongside detailed characterisation of vegetation structure and ground□layer microhabitats. In total, 38 species and 25,578 individuals were recorded. Microhabitat variables explained 58% of the variation in species occurrence. Community differences among microhabitats were driven primarily by nestedness, with dense herbaceous cover acting as a core habitat and edge□related components contributing disproportionately to beta diversity. Tree abundance showed a unimodal relationship with species richness, with maximum diversity at intermediate densities, while shrub and lawn cover had weak or inconsistent effects. Fine□scale elements such as leaf litter, stones, woody debris, and small bare□ground patches strongly influenced species occurrence by providing thermal refugia, nesting substrates, and foraging opportunities. The invasive Argentine ant (*Linepithema humile*) exhibited strong but spatially restricted dominance and species□specific negative effects on native ants, emphasising the role of habitat context in mediating invasion impacts. Our results demonstrate that urban biodiversity is maximised by enhancing fine□scale habitat heterogeneity rather than increasing green cover alone. We highlight practical design principles for urban green infrastructure that prioritise structural diversity and ground□layer complexity to support resilient arthropod communities.

## 1. Introduction

Global change has been identified as one of the major drivers of biodiversity loss (Jaureguiberry et al., 2022; United Nations Environment Programme - UNEP, 2023). As a defining feature of the Anthropocene, it encompasses a suite of environmental transformations driven by human activities, including land and sea use change, pollution, climate change, overexploitation of organisms (i.e. overfishing, hunting, wildlife trade…) or biological invasions (Brondízio et al., 2019; Maxwell et al., 2016; Niederman et al., 2025). Collectively, these transformations are exerting profound impacts on wildlife and human health (Díaz et al., 2019) by degrading ecosystem structure, functioning, and resilience; ultimately diminishing the environmental quality upon which life depends (*Connecting Global Priorities*, 2015). Among all drivers of global change, urbanization deserves special attention, challenging organisms survival through warmer temperatures (Dietzel et al., 2025; Jabbar et al., 2023), landscape shifts and fragmentation (Faeth et al., 2011; Marzluff, 2008; Sattler et al., 2010), increase of artificial light at night (Miller et al., 2017; Tougeron & Sanders, 2023) or soil and air pollution (Badia et al., 2025; Hanif & Benitez, 2025; Moura et al., 2025), and the establishment of invasive alien species (Bertelsmeier, 2021; Padayachee et al., 2017; Shochat et al., 2010; Trentanovi et al., 2013). While some organisms may thrive in these artificial, novel environments, others face far more severe outcomes (Andersen, 2019; Dunn et al., 2022; Sanllorente et al., 2025; Sattler et al., 2010), including population declines or even local extinctions.

Both in plants and animals, the impact of the urban pressures is stronger depending on their ecological plasticity and movement capacity (Hahs et al., 2023; Ruas et al., 2022; Sexton et al., 2025). In plants, dispersal ability largely depends on the efficiency of seed dispersal mechanisms, which determine the capacity of species to colonize or persist in fragmented habitats (Mendes et al., 2025; Nathan & Muller-Landau, 2000). In animals, dispersal capacity – particularly that of females, which often drive population growth and colonization dynamics – can strongly influence population persistence. This limitation is especially relevant in species with gregarious or colony based social systems, such as ants, where population viability depends on a small number of reproductive individuals (King & Tschinkel, 2016).

Ants are widespread organisms present in almost all habitat types, although community composition varies depending on different environmental factors (Schultheiss et al., 2022). Broad-scale climatic patterns – operating at both large and small spatial scales (i.e., biogeographic regions and local climatic conditions, respectively) – are major determinants of ant diversity and largely define the potential myrmecofauna capable of colonizing a region (Dantas & Fonseca, 2023; Tinaut & Ruano, 2021). For instance, the general climatic regime of a region (e.g., tropical versus Palearctic) sets the regional species pool. However, at finer spatial scales, microhabitat characteristics ultimately determine the realized ant diversity at a site, as well as the composition of their associated fauna (Barton et al., 2024; Parmentier & Braem, 2023; Queiroz et al., 2013; Tenorio-Escandón et al., 2023; Trigos-Peral et al., 2018), according to the ecological requirements of the occurring species (Seifert, 2017).

This fine □scale dependence on microhabitat structure is particularly relevant in urban green spaces, where simplified habitat configuration can act as a strong environmental filter. Such simplification may constrain the availability of nesting sites and resources, potentially increasing species turnover among sites (Dijon et al., 2023; Vepsäläinen et al., 2008; Yin et al., 2024) by constraining the availability of nesting sites and resources. Moreover, some components of the microhabitat also serve as refuge for species in harsh environments, providing protection towards high temperatures (Scheffers et al., 2014). This buffering function is especially important in southern Mediterranean regions, where ants are exposed to extremely high surface temperatures during the warmest months, frequently exceeding 50–60□°C during peak insolation on bare soil (Cerdá et al., 1998; Illán-Fernández et al., 2025; Proutsos et al., 2025).

In addition to habitat simplification, urban native fauna, including ants, face increasing pressure from invasive species. Among them, the Argentine ant, *L. humile*, is the most widespread invasive ant species in the Mediterranean region. Beyond its substantial economic impacts (Angulo et al., 2021), its invasion success is favoured by warm climatic conditions (Abril, 2010; Abril, Oliveras, et al., 2010) and the predominance of anthropogenic habitats (Angulo et al., 2024; Calcaterra et al., 2025; Roura-Pascual et al., 2011). Moreover, its large polydomous and polygynous colonies efficiently monopolize food and nesting resources, restricting native ants’ access to these key resources through direct competition and behavioural exclusion (Abril & Gómez, 2011; Buczkowski & Bennett, 2008; Holway, 1999; Holway et al., 2002; Human & Gordon, 1996; Madrzyk et al., 2026). Due to their broad ecological and behavioural plasticity, as well as strong numerical and hierarchical dominance (Bertelsmeier et al., 2016; Blight et al., 2017; Espadaler & Rey, 2001; Trigos-Peral et al., 2021; Muñoz et al., 2026), invasive ants can substantially reduce the persistence of native species in highly modified urban habitats, frequently leading to severe population declines or local collapse. These impacts highlight the importance of integrating biodiversity-sensitive approaches into urban greenery design. In this context, urban green infrastructure is increasingly recognised not only as a provider of ecosystem services but also as a key component for supporting native biodiversity within highly modified landscapes, reinforcing its role as urban refuges for fauna (Capotorti et al., 2017; Perrelet et al., 2024). This perspective promotes the incorporation of habitat features that enhance the persistence of native species while potentially reducing the ecological advantages of invasive species.

Given the importance of carefully selecting key microhabitat components in the design of urban green spaces to enhance their function as refuges for native biodiversity, particularly during periods of harsh climatic conditions such as summer, this study evaluates how these structural elements relate to ant occurrence in urban green spaces. Specifically, we examine whether ant distributions are randomly organised or instead associated with distinct microhabitat components shaped by landscape design and management practices. In addition, the presence of the Argentine ant, *Linepithema humile*, in part of the study area allows us to assess whether this invasive species disproportionately exploits key microhabitat resources, with potential implications for designing urban greenery configurations that support native ant assemblages while limiting competitive dominance by invasive species.

## 2. Material and Methods

### 2.1. Study areas and sampling method

Sampling was conducted across twelve urban green spaces at different locations in the city of Cordoba (Spain), including public parks, a botanical garden, and university gardens. The study consisted in periodic yearly sampling over a period of 10 years (2004–2013, both included) during the months of June or July, matching with the peak ant activity in the south of the Iberian Peninsula. As sampling methodology, we used transects with 10 pitfall traps separated by 5 m distance. In each green space, we placed 2 or 3 transects – depending on its size (Table 1) – separated by a minimum distance of 20 m.

**Table 1.**
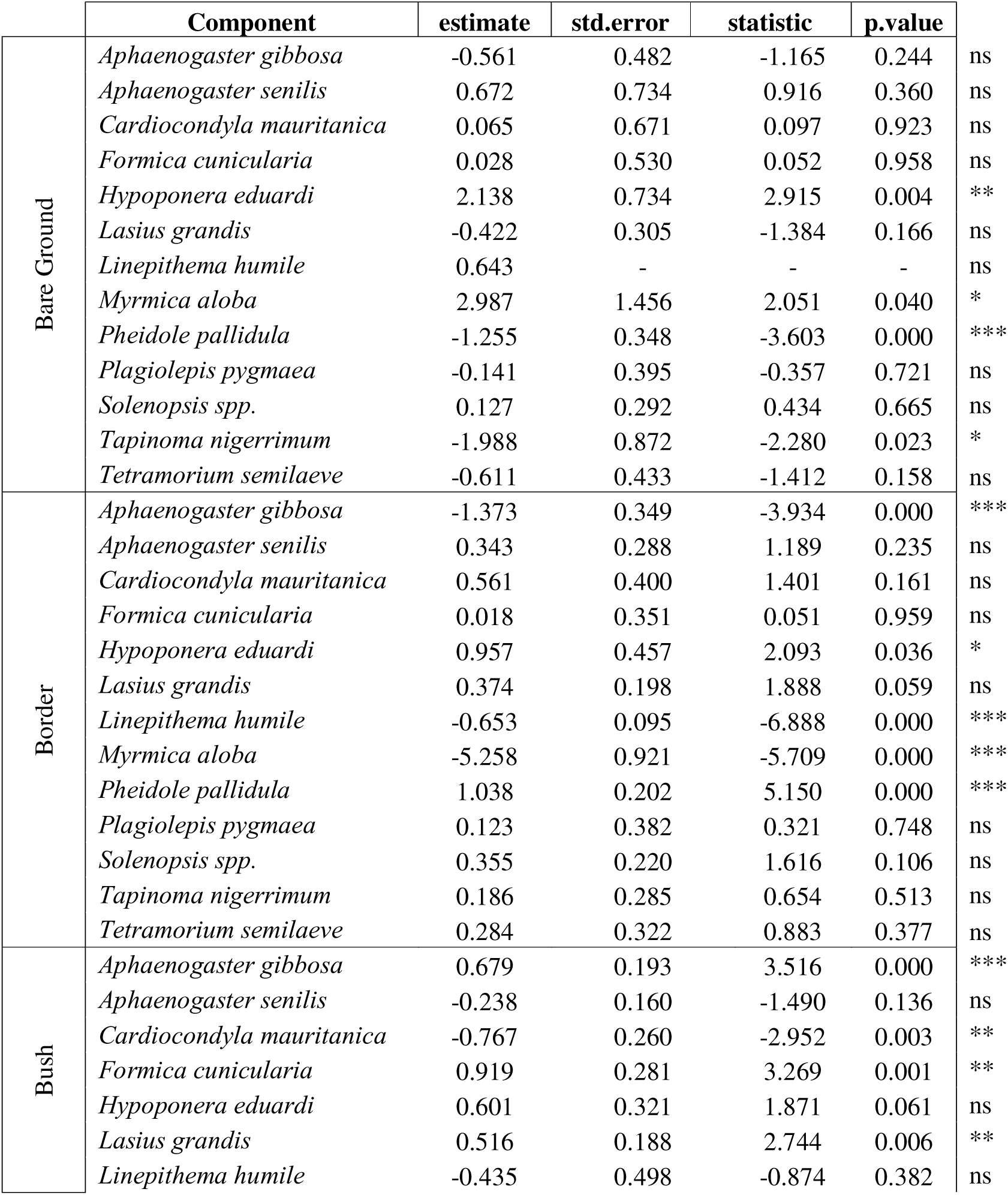
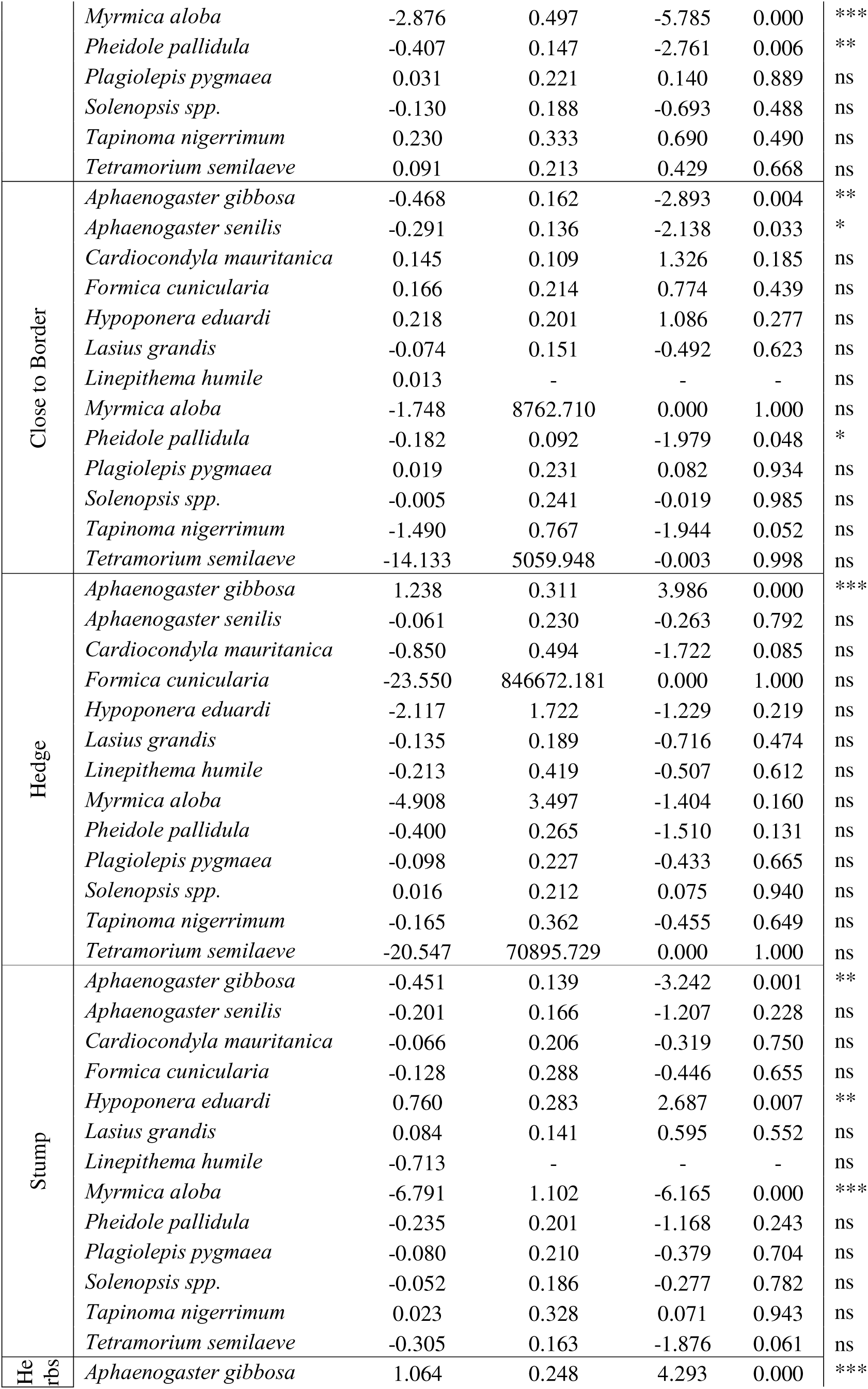
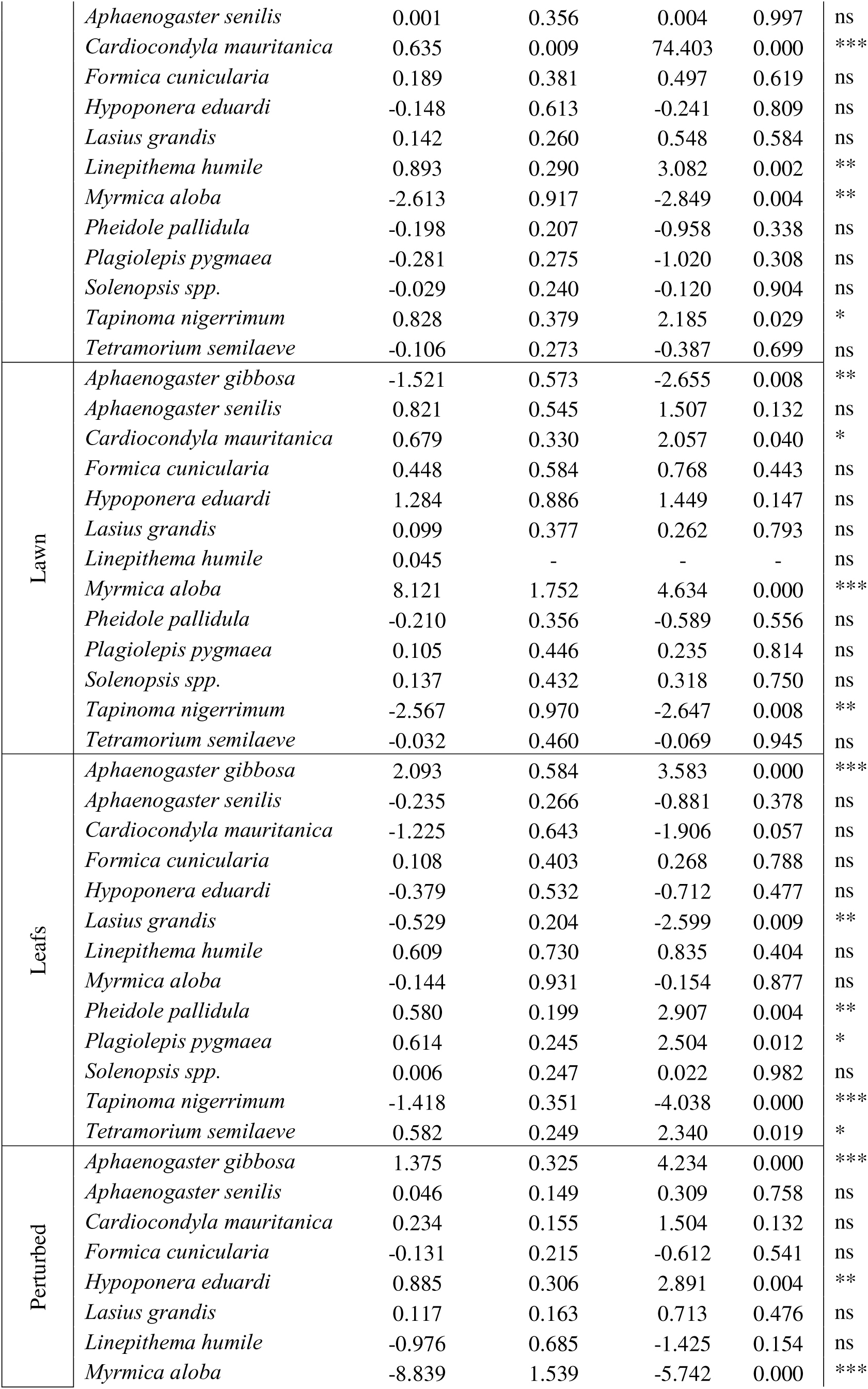
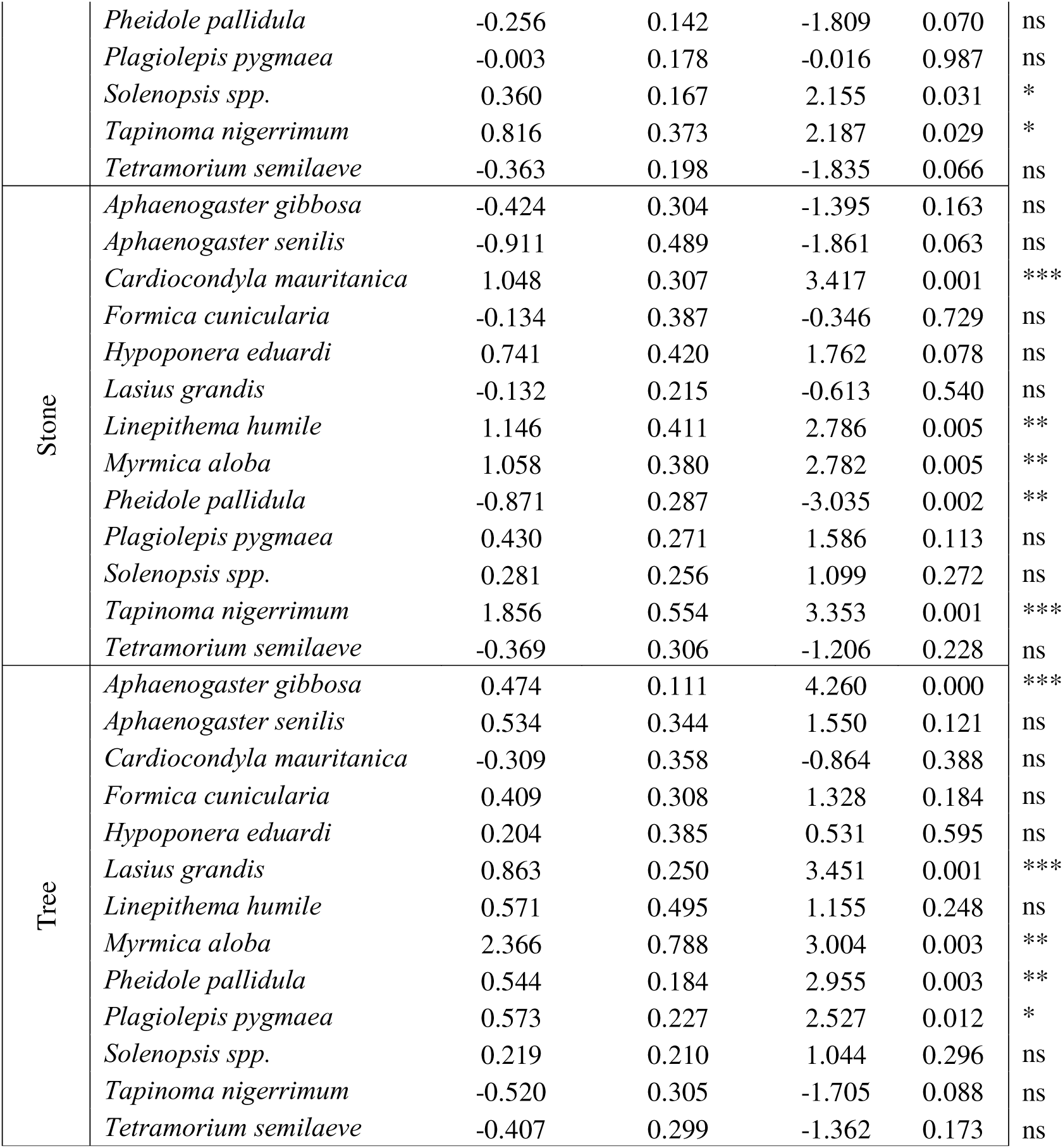
Results of the generalized linear mixed models (Negative Binomial distribution) examining the relationship between each microhabitat component and the occurrence of the most representative ant species in urban green spaces. Non-significant effects are indicated as “ns”, whereas significant effects are denoted by asterisks (*).

We used semi-transparent plastic cups (7.3 cm deep, 5.7 cm top diameter, 5 cm base; REF. 409702, DELTALAB SL) as pitfall traps, filled with 50 ml of water and a drop of detergent to prevent ants from escaping. Traps were placed in the soil for 48 h, after which collected specimens were sorted and identified to species under a stereomicroscope.

To inventory the microhabitat components, we photographed each pitfall trap along with its surrounding area. The photos were then examined to identify the microhabitat features around each trap, resulting in the identification of 16 distinct parameters: Tree foot, Under tree, Bush foot, Under bush, Close to stump, Close to hedge, Close to border, Grass, Dense grass, Bare ground, Herbaceous, Dense herbaceous, Stone, Leafs, Perturbed, Border (Supplementary table 1). To avoid collinearity during some statistical analyses, microhabitat components were grouped into broader categories: *Tree* (tree foot and under tree), *Bush* (bush foot and under bush), *Herbs* (herbaceous and dense herbaceous) and *Borders* (close to border and border). In addition, we defined four major categories based on general similarities in habitat characteristics: *Wood* (indicating the presence of trees, bushes, or stumps), *Green cover* (referring to areas where the soil surface is densely covered by grass, herbaceous vegetation, leaf litter, or hedges), *Open* (describing areas with direct insolation, such as bare ground, perturbed soil, or sparsely vegetated patches) and *Habitat shift* (encompassing microhabitats characterized by abrupt changes in structure or composition, such as borders or the presence of stones).

### 2.2. Statistical analyses

All statistical analyses were carried out with R (R Core Team 2020). We created graphical representations of the results using the *ggplot* function in the *ggplot2* package. Species richness, species turnover (Jaccard similarity index and nestedness, and Mantel test), Detrended Correspndence Analyses (DCA) and Canonical Correspondence Analyses (CCA) were carried out using the *vegan* package (Oksanen et al., 2025). Generalized Linear Mixed Models (GLMMs) were carried out using the *glmmTMB* package (McGillycuddy et al., 2025). Quadratic GLMs were carried out using the *MASS* package (Venables & Ripley, 2009). All GLMMs were tested for model fit using *DHARMa* package (Hartig et al., 2025). Plots were created using the *ggplot2* package (Wickham, 2009).

First, the multivariate approach was selected by performing a DCA on the matrix of species workers abundance, which Axis length value (length > 4) supported the use of CCA. To check whether the different components play a role in the ant species exploiting the microhabitat, we performed a CCA on the matrix of number of workers present in the immediate location of each component (as the number of workers of each ant species in pitfall traps) and the presence of each component. Later, we carried out an Anova on the CCA model to check for the significance of each component. We use the raw number of workers as both the number of foragers or the presence of a colony in this exact location is directly connected to the suitability of the microhabitat to the species.

To examine species turnover and nestedness among the microhabitat components, we calculated the Jaccard similarity index and nestedness on the matrix of the presence/absence of species and microhabitat components. Later, we calculated the Mantel test on the Jaccard-based turnover matrix to test whether the species turnover is structured by these components in a distance-based way. For species turnover, values were interpreted as follows: < 0.1 – low species turnover, 0.1-0.3 – medium species turnover, >0.3 - high species turnover.

Later, we evaluated the optimal density of the three key microhabitat components commonly used in urban greenery design in relation to ant species richness. To this end, we applied quadratic generalized linear models (Negative Binomial distribution) based on the formula

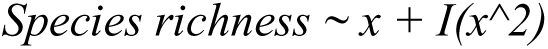

in which x estimates the linear effect of the microhabitat components and I(x^2) estimates their quadratic effect. This analysis was applied to the abundance matrix of workers of the numerically dominant species and the abundance of the following habitat components: trees (tree foot + under tree), shrubs (bush foot + under bush), and lawns (grass + dense grass). To assess whether additional microhabitat components could be relevant for urban greenery design aimed at enhancing the role of urban green spaces as refuges for biodiversity, we fitted zero-inflated generalized linear mixed models (GLMMs; family = nbinom2). The response variable was the abundance of workers for each species, while the explanatory variables were the abundances of individual microhabitat components per transect. In the models, the identity of the garden and the transect were used as nested random factor. To simplify the graphical visualization of the relationship between microhabitats and species occurrence, we used the matrix of abundance of the main native ant species and presence/absence of the four main microhabitats categories: *Wood*, *Green Cover*, *Open* and *Habitat shift*.

We used worker abundance as a proxy for microhabitat suitability, assuming that higher numbers of workers reflect either greater colony success or a stronger preference for a given location during foraging. Because sampling was conducted in late spring and summer – periods characterized by extremely high temperatures in the southern Iberian Peninsula – higher worker abundance is likely to provide a reliable indication of the relative advantages associated with each microhabitat component. To reduce statistical noise, analyses were restricted to species recorded in at least 3% of traps and represented by more than 100 workers.

Finally, to check whether the presence of the invasive Argentine ant (*L. humile*) can interfere in the use of these microhabitat components by native species, we performed GLMMs (Negative binomial distribution). In these models, species abundance was the response variable, while the presence/absence of Argentine ant per trap served as the explanatory variable. Transect and trap identity were included as nested random effects. For these analyses, we only use data from those parks in which the Argentine ant was present.

## 3. Results

Our results are originated from the data collected during the 10-year period. In total, we analysed ant diversity from 681 pitfall traps collected in 11 urban parks in Córdoba (Spain). A total of 38 species were identified from 25,578 individuals.

Partitioning Jaccard dissimilarity revealed that community differences among microhabitats were dominated by nestedness, with most assemblages representing subsets of a species-rich dense herbaceous core habitat. Grass, dense grass, herbaceous vegetation, trees, bushes, leaves, stone, and bare ground exhibited moderate to high nestedness, indicating compositional differences driven primarily by species loss rather than replacement. In contrast, edge-related microhabitats (close hedge, close border, and close stump) showed lower nestedness coupled with higher turnover, reflecting stronger species replacement and ecological filtering at habitat boundaries. Tree and bush microhabitats displayed consistently high nestedness and low turnover with surrounding habitats, suggesting limited contribution to beta diversity and a largely shared species pool. Overall, dense herbaceous vegetation acted as a central source habitat, while edge microhabitats disproportionately contributed to beta diversity through species turnover rather than nestedness (Fig. 1, Supplementary Table 1, Supplementary Figure 1).

**Figure 1.**
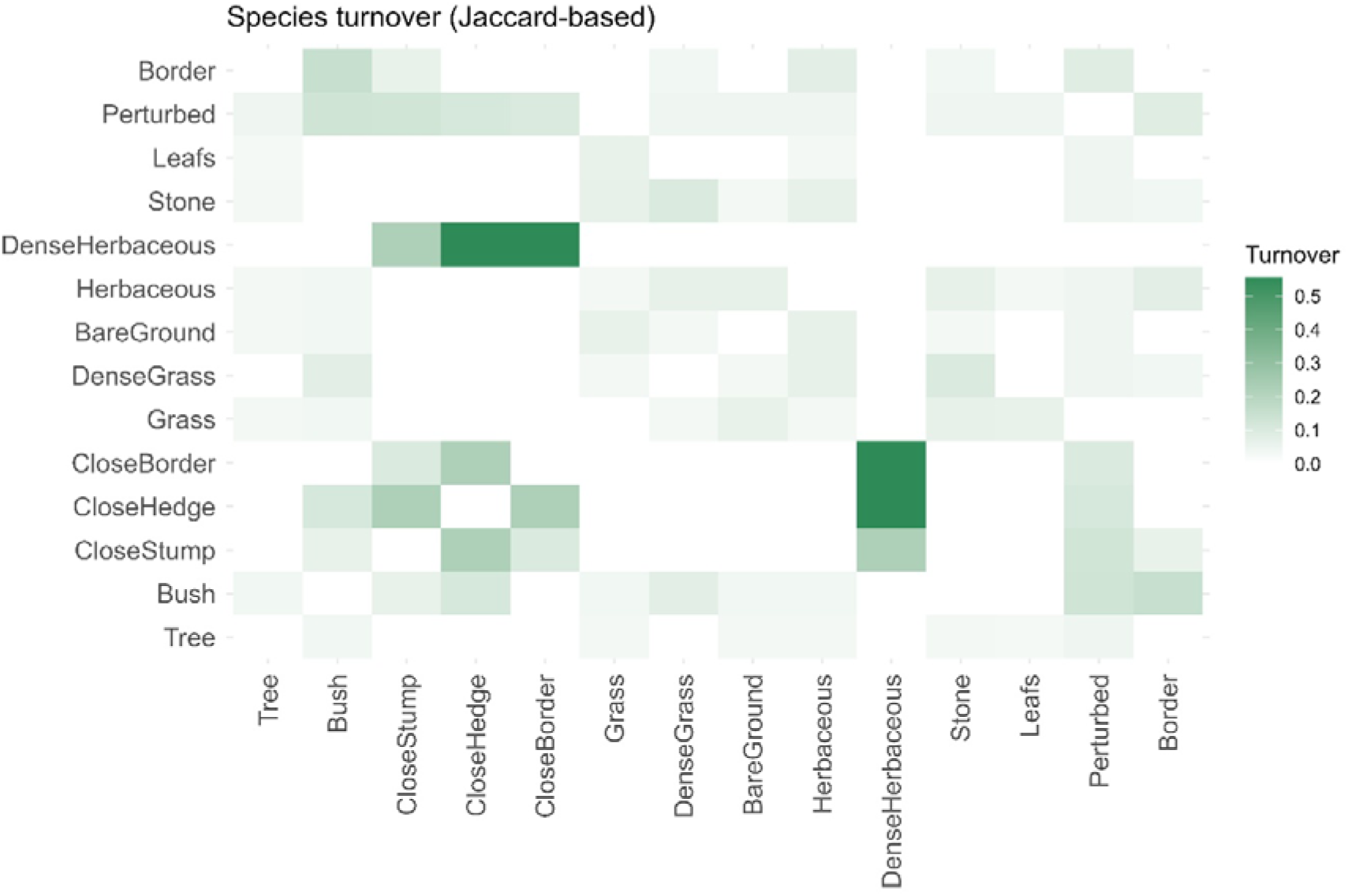
Heatmap showing Jaccard-based species turnover among the different components of the microhabitat. Higher turnover is indicated by darker green squares.

Results of the CCA indicate that microhabitat components explain 58.34% of the variation in ant species occurrence. The first two ordination axes accounted for 34.65% (CCA1) and 23.69% (CCA2) of the variation in species abundance, respectively (Figure 2). Moreover, these components showed a significant influence on the species composition (F = 6.37, df = 13, p = 0.001), with the strongest influence found with the presence of woody vegetation (Tree: F = 12.65, p = 0.001; Bush: F = 13.39, p = 0.005), the coverage of the ground (Grass: F = 6.72, p = 0.02; Dense grass: F = 8.56, p = 0.05; Herbaceous: F = 5.35, p = 0.33; Leafs: F = 11.15, p = 0.003) and the presence of medium/large size stones (F = 6.16, p = 0.027).

**Figure 2.**
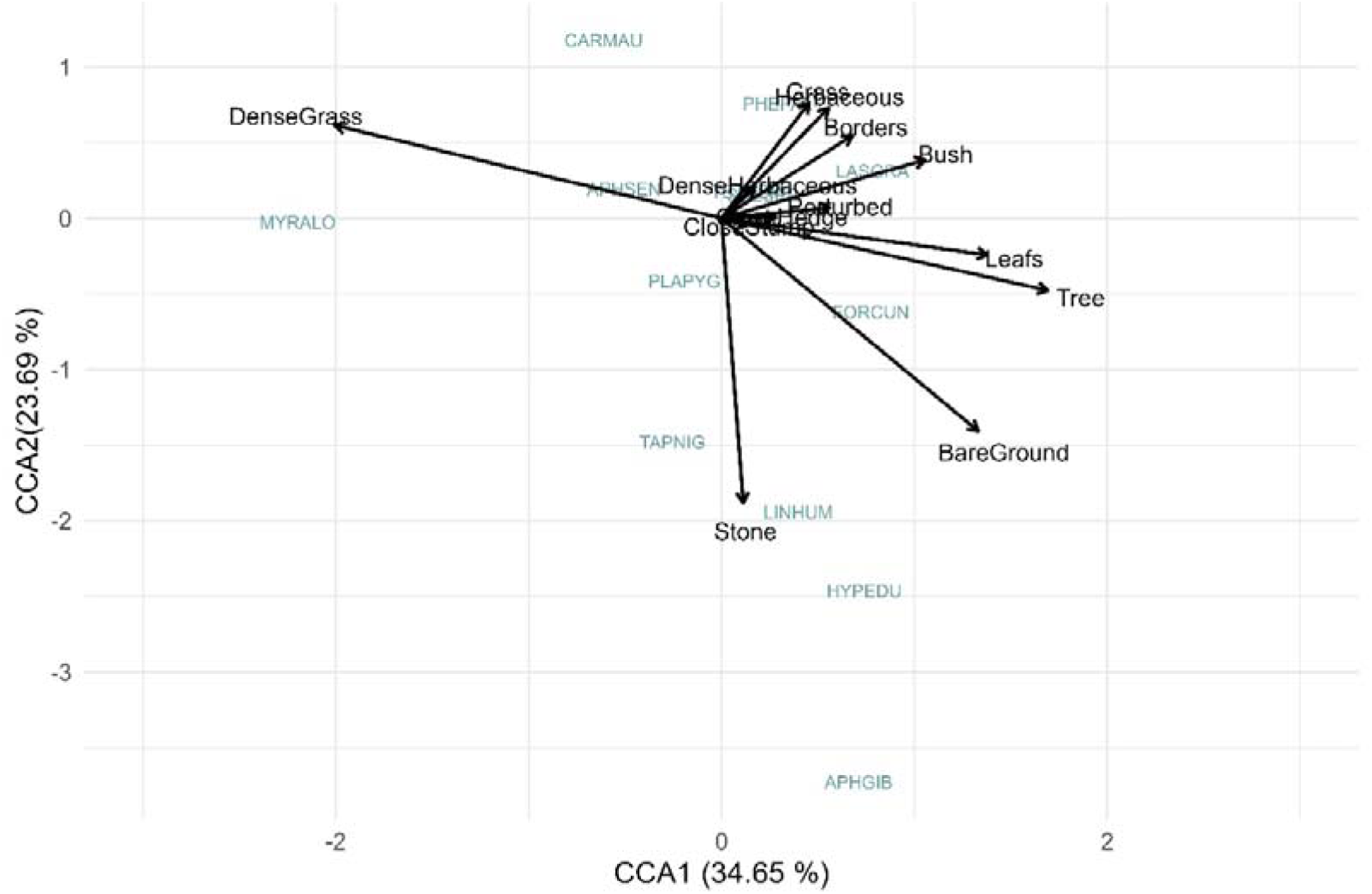
Canonical correspondence analysis (CCA) biplot showing the relationships between species composition and microhabitat components.

The use of the main microhabitat components in urban parks design showed to have a different impact on the biodiversity. Species richness was significantly related to tree abundance, showing a nonlinear response, whereas shrub and lawn abundance had no significant effects. The relationship with tree abundance was hump-shaped, indicating higher species richness at intermediate tree abundance and lower values at both low and high levels. Tree abundance showed a positive linear effect (β = 0.173, SE = 0.084, p = 0.040) together with a significant negative quadratic effect (β = −0.018, SE = 0.009, p = 0.044), confirming this unimodal pattern. In contrast, shrub abundance (linear: β = −0.045, SE = 0.076, p = 0.553; quadratic: β = 0.012, SE = 0.014, p = 0.387) and lawn abundance (linear: β = 0.036, SE = 0.040, p = 0.365; quadratic: β = −0.002, SE = 0.002, p = 0.472) did not significantly influence species richness (Figure 3).

**Figure 3.**
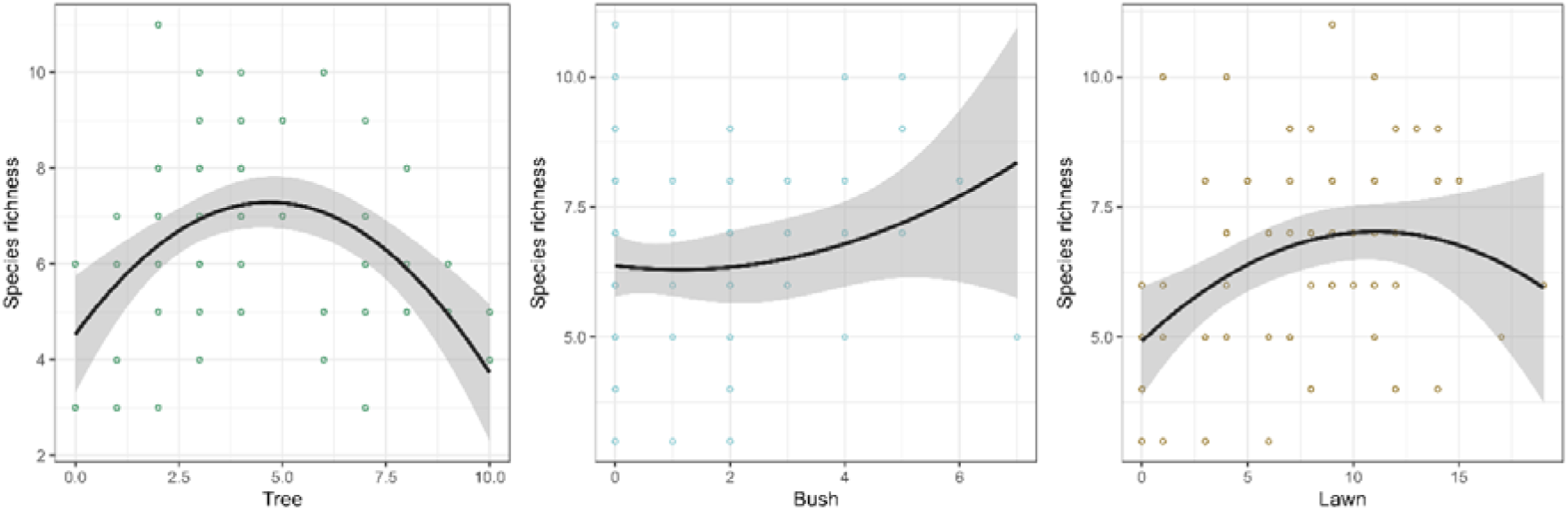
Relationship between ant species richness and the abundance of main microhabitat components in urban green spaces: trees (left), bushes (centre), and lawns (right). Points represent observed values per sampling unit, while solid lines show fitted predictions from quadratic generalized linear models. Shaded areas indicate 95% confidence intervals.

When evaluating which microhabitat components are most relevant for biodiversity-oriented urban green-space design, we found that structurally complex microhabitats, especially those combining arboreal cover and litter accumulation, play a central role in promoting ant diversity in urban green spaces, whereas simplified or highly exposed habitats support a more limited and selective assemblage of species (Table 1, Figure 4).

**Figure 4.**
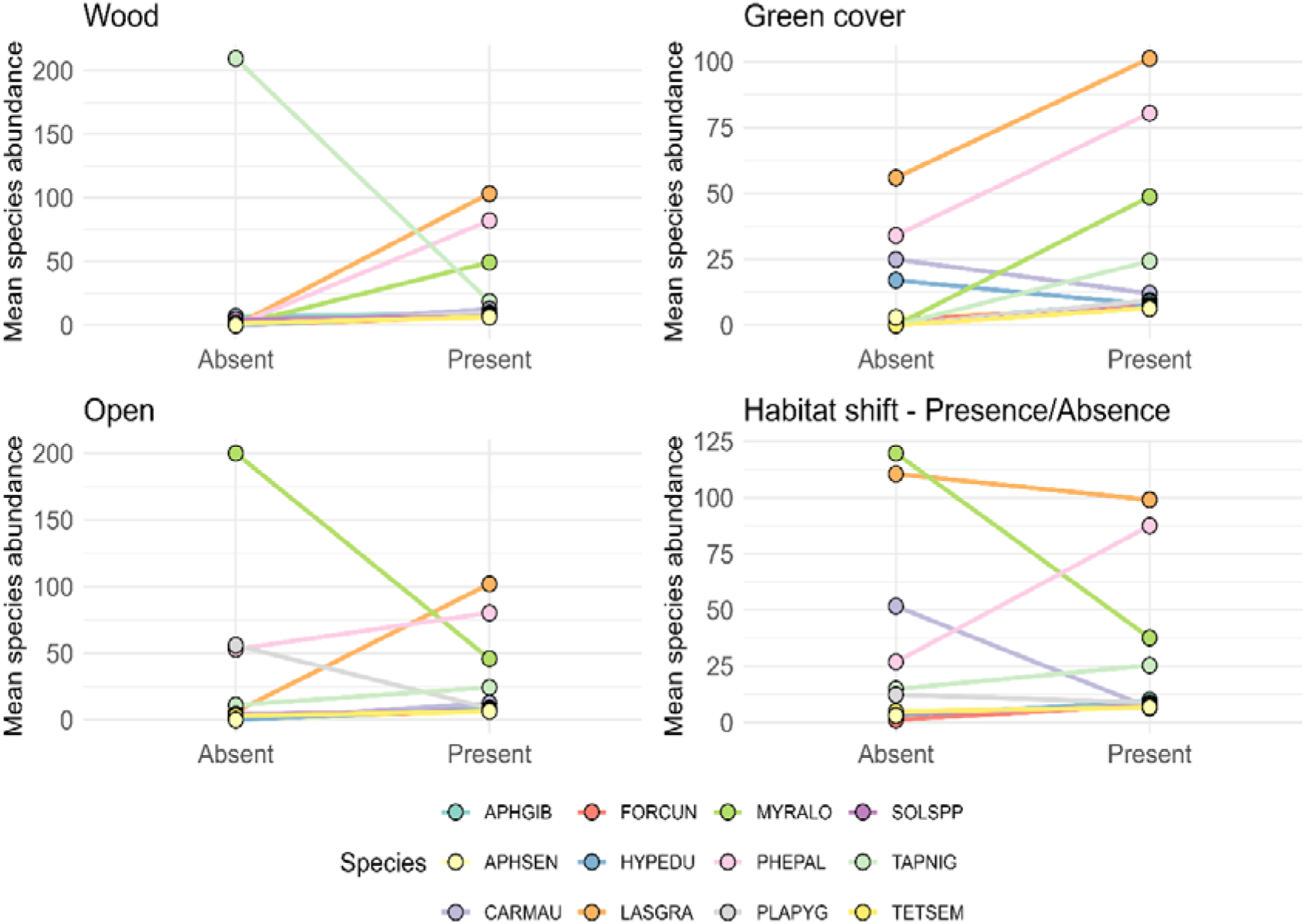
Graphical representation of the variation on the species abundance at microhabitat level under presence and absence of different microhabitat components: Wooden components (trees, bushes and stumps), green cover (leaflitter, lawns, dense herbaceous and hedges, Open spots (sparse grass cover, perturbed surface and bare ground) and Habitat shift (border areas and stones). Abbreviations correspond to: *Aphaenogaster gibbosa* (APHGIB), *Aphaenogaster senilis* (APHSEN), *Cardiocondyla mauritanica* (CARMAU), *Formica cunicularia* (FORCUN), *Hypoponera eduardi* (HYPEDU), *Lasius grandis* (LASGRA), *Myrmica aloba* (MYRALO), *Pheidole pallidula* (PHEPAL), *Plagiolepis pygmaea* (PLAPYG), *Solenopsis* spp. (SOLSPP), *Tapinoma nigerrimum* (TAPNIG), *Tetramorium semilaeve* (TETSEM).

More specifically, microhabitats associated with tree presence are the most species-rich, showing positive associations with 10 of the 13 studied species. This effect is statistically significant for five species, highlighting the strong contribution of tree-related microhabitats to overall species occurrence patterns. Similarly, leaf-litter microhabitats supported a high number of species, with positive associations observed for eight species and significant effects detected for *A. gibbosa*, *P. pallidula*, *P. pygmaea*, and *T. semilaeve*, whereas *L. grandis* and *T. nigerrimum* showed significant negative associations. Herbaceous vegetation and perturbed microhabitats also exhibited relatively high species richness, each being positively associated with eight species, although the strength and significance of these relationships differed among taxa. In contrast, bare ground and lawn habitats showed fewer positive associations and were characterized by stronger species-specific responses, including significant negative associations for several dominant taxa such as *P. pallidula*, *T. nigerrimum*, and *A. gibbosa*. Stone-associated microhabitats produced contrasting responses, positively influencing disturbance-tolerant species such as *C. mauritanica*, *L. humile*, and *T. nigerrimum*, while negatively affecting *P. pallidula*. Border and close-border microhabitats were generally associated with fewer species and frequently showed negative relationships, particularly for *A. gibbosa*, *L. humile*, and *M. aloba* (Table 1, Figure 4).

### Presence of Argentine ant and monopolization of resources

The presence of *L. humile* was restricted to three of the twelve studied green spaces. Within these sites, this invasive species exhibited marked variation in trap occupancy, accounting for 89.2% of the collected traps in the Botanical Garden, followed by 25.4% in CM-Asunción, and reaching its lowest level in Jardines de Renfe with 15% occupancy.

Overall, the impact of *L. humile* within urban green spaces appeared to be species-specific, disproportionately affecting dominant native taxa while exerting weaker or negligible effects on others. Significant negative associations were detected for several of the most common species inhabiting the studied green spaces, suggesting competitive exclusion or reduced access to resources in sites occupied by the invasive species. In contrast, *Myrmica aloba* showed a significant positive association with *L. humile*. No significant relationships were observed for *Cardiocondyla mauritanica*, *Plagiolepis pygmaea*, *Aphaenogaster gibbosa*, *Solenopsis* spp., or *Aphaenogaster senilis* (Table 2, Fig. 5).

**Figure 5.**
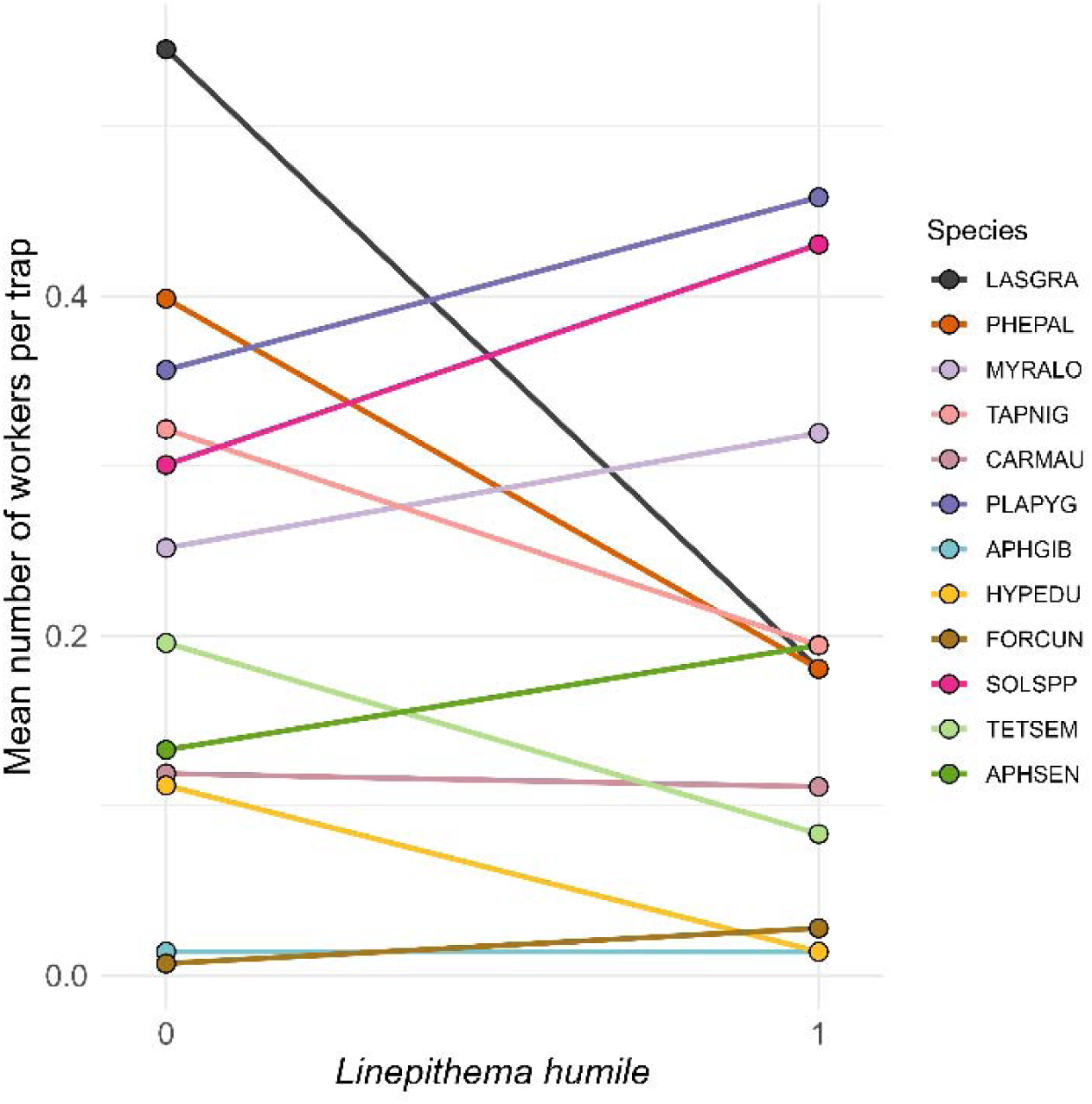
Graphical representation of the mean number of workers per trap for the most representative ant species in traps where *Linepithema humile* was absent (0) or present (1). Values represent the mean worker abundance per trap across the three invaded urban green spaces (Botanical garden, CM-Asuncion, Jardines de Renfe).

**Table 2.** Species-specific associations between ant occurrence and urban microhabitat components obtained from generalized linear mixed models.

| Species | estimate | std.error | statistic | p.value | p.adj<br>holm |  |
| --- | --- | --- | --- | --- | --- | --- |
| *Lasius grandis* | -2.362 | 0.107 | -21.993 | < 0.001 | < 0.001 | *** |
| *Pheidole pallidula* | -0.818 | 0.112 | -7.325 | < 0.001 | < 0.001 | *** |
| *Myrmica aloba* | 0.550 | 0.036 | 15.411 | < 0.001 | < 0.001 | *** |
| *Tapinoma nigerrimum* | -3.116 | 0.225 | -13.835 | < 0.001 | < 0.001 | *** |
| *Cardiocondyla mauritanica* | -0.427 | 0.344 | -1.240 | 0.215 | 0.645 | ns |
| *Plagiolepis pygmaea* | 0.258 | 0.122 | 2.111 | 0.035 | 0.174 | ns |
| *Aphaenogaster gibbosa* | 1.973 | 2.287 | 0.863 | 0.388 | 0.753 | ns |
| *Hypoponera eduardi* | -3.157 | 1.021 | -3.093 | 0.002 | 0.012 | * |
| *Solenopsis* spp. | 0.139 | 0.157 | 0.884 | 0.377 | 0.753 | ns |
| *Tetramorium semilaeve* | -1.135 | 0.236 | -4.812 | < 0.001 | < 0.001 | *** |
| *Aphaenogaster senilis* | 0.295 | 0.206 | 1.434 | 0.152 | 0.606 | ns |

## 4. Discussion

The design and management of ecologically sustainable urban green spaces have become central strategies for mitigating biodiversity loss in increasingly urbanised landscapes. Traditionally, urban green-space planning was largely driven by aesthetic and recreational considerations, often prioritising simplified vegetation structures and intensive maintenance regimes. In recent decades, however, urban landscape design has progressively shifted towards more multifunctional approaches that integrate visual attractiveness with ecological functionality and biodiversity conservation objectives. This transition has promoted the incorporation of structurally diverse vegetation, native plant assemblages, habitat heterogeneity, and microhabitat features capable of supporting a wider range of native species while simultaneously enhancing ecosystem services and human well-being. As a result, urban green spaces are increasingly recognised not only as recreational infrastructures but also as critical components of urban biodiversity conservation networks and ecological resilience strategies (Bryant, 2006; Hoyle et al., 2017; Heymans et al., 2019; Kowarik et al., 2025).

In parallel, a growing body of literature has documented the effects of urbanisation-related stressors on biodiversity across taxa (Davis & Glick, 1978; Shochat et al., 2010; Lososová et al., 2012; Trentanovi et al., 2013; Sanllorente et al., 2025; among others). Given their ecological importance and sensitivity to environmental change, arthropods – particularly ants – have received considerable attention (Vepsäläinen et al., 2008; Sattler et al., 2010; Ślipiński et al., 2013; Carpintero & Reyes-López, 2014; Trigos-Peral et al., 2020; Trentanovi et al., 2013; Ferrari et al., 2025; Trigos-Peral, Maák, et al., 2024; Trigos-Peral & Reyes-López, 2025). Ants play a fundamental role in ecosystem functioning, contributing to processes such as soil turnover, seed dispersal, and trophic regulation (Gardiner et al., 2013; Kaiser & Resasco, 2024). However, despite increasing empirical evidence on urban impacts, the underlying mechanisms driving changes in ant community structure remain insufficiently understood (Lokatis & Jeschke, 2022; Penick & Edenborough, 2026). In particular, fine-scale studies addressing microhabitat-level processes remain scarce (Queiroz et al., 2013; Parmentier & Braem, 2023), despite their importance in providing food resources, nesting sites, and thermal refugia under harsh urban conditions (Angiletta, 2010; Diamond et al., 2017).

### 4.1. Vegetation structures as key microhabitat components for native species conservation

Our results support previous findings (Barton et al., 2024), demonstrating that microhabitat structure is a major driver of ant community composition in urban green spaces. Among the variables considered, tree abundance emerged as the most influential factor shaping species richness and assemblage structure. This pattern highlights the importance of vegetation architecture in urban design, particularly given the role of tree canopies in moderating microclimatic extremes by reducing surface temperatures, increasing soil moisture retention, and providing shaded refuges during warm periods (McDonnell et al., 2011; Wenzel et al., 2020; Beaumelle et al., 2021).

Notably, the relationship between tree abundance and ant diversity was non-linear, with intermediate levels of tree cover maximising species richness. This hump-shaped response is consistent with the intermediate disturbance and structural complexity hypotheses, whereby biodiversity peaks at moderate levels of habitat complexity or environmental stress (Connell, 1978; Arnan et al., 2017). Both simplified systems with low tree density and overly closed canopies may reduce habitat suitability: open areas are exposed to thermal and desiccation stress, while dense canopies may limit understorey development and ground-level heterogeneity critical for ant foraging and nesting (Gibb & Hochuli, 2003; Maák et al., 2021). Species-level responses further illustrate these mechanisms. Meadows and litter-dependent taxa such as *A. gibbosa*, *L. grandis*, *M. aloba*, and *P. pallidula* showed strong positive associations with tree presence, while thermophilous or disturbance-tolerant species responded weakly or negatively. Similar species-specific tree effects have been documented across urban and semi-natural systems, highlighting the role of canopy cover in filtering ant assemblages through microclimatic and resource-mediated pathways (Yamaguchi, 2004; Buczkowski & Richmond, 2012).

In contrast, shrub and lawn cover exhibited weak or inconsistent effects on overall ant diversity. Although these elements are commonly incorporated into urban planting schemes, their ecological contribution appears to depend more on spatial configuration, connectivity, and integration within a heterogeneous matrix than on proportional cover alone (Beninde et al., 2015; Threlfall et al., 2017). Lawns, in particular, represent structurally homogeneous habitats subjected to frequent mowing and soil compaction, conditions known to reduce arthropod diversity and favour a limited subset of disturbance-tolerant taxa (Azcárate & Peco, 2011). However, mosaic of scattered and dense grass areas in the studied green spaces have shown to substantially dump its impact, or even favouring the presence of both hygrophilous species (*M. aloba*) and more thermophilic ones with preference for open spots (*C. mauritanica*).

### 4.2. Role of litter and other fine-scale habitat complexity

Beyond vegetation structure, additional microhabitat components such as leaf litter, bare ground, stones, and fine-scale disturbance play a critical role in shaping ant communities. Forest structure and litter accumulation generate a mosaic of microhabitats that provide shelter, foraging opportunities, and nesting sites for ants (Levings, 1983; Yanoviak & Kaspari, 2000). In urban and semi-natural systems alike, leaf litter influences soil arthropod communities by moderating microclimatic conditions, buffering temperature and moisture fluctuations, and regulating nutrient cycling processes (Bradford et al., 2002; Vasconcelos & Laurance, 2005).

In our study, leaf-litter presence positively affected several native and litter-specialist species, including *A. gibbosa*, *P. pallidula*, *P. pygmaea*, and *T. semilaeve*. These patterns align with observations from tropical and temperate ecosystems, where litter depth and heterogeneity are positively correlated with ant abundance and diversity (Kaspari & Yanoviak, 2009; Poeydebat et al., 2021). By contrast, although intensive litter removal is a common practice in urban parks, it can severely limit nesting opportunities and increase exposure to thermal stress, leading to simplified assemblages dominated by disturbance-tolerant species (Magura & Lövei, 2021). Bare ground and lawn habitats showed more selective effects. Exposed substrates favoured a limited number of species tolerant to extreme temperatures and low moisture, such as *H. eduardi*, but were negatively associated with more mesophilous taxa. Similarly, lawns promoted high abundances of *M. aloba* while reducing forest-associated taxa such as *A. gibbosa*, supporting previous evidence that homogeneous turfgrass systems act as ecological filters in urban landscapes (Norton et al., 2019). Stones and woody debris further contributed to fine-scale heterogeneity, benefiting species capable of exploiting crevices and thermal refugia, including *C. mauritanica*, *Linepithema humile*, and *T. nigerrimum*. Such elements can substantially increase local diversity when embedded within heterogeneous matrices, despite occasionally favouring opportunistic or invasive species (Muñoz et al., 2026b).

### 4.3. Edge effects, disturbed habitats, and urban-tolerant assemblages

Edge-related habitats (e.g. borders, hedges, and transition zones) produced pronounced and species-specific responses. Borders negatively affected forest-associated taxa such as *A. gibbosa* and *M. aloba*, while positively influencing generalist species like *P. pallidula*. These patterns are consistent with edge effects documented in fragmented and urbanised landscapes, where increased temperature, reduced humidity, and frequent disturbance favour opportunistic and competitively dominant species. Perturbed habitats promoted assemblages dominated by disturbance-tolerant species, including *H. eduardi*, *Solenopsis* spp., and *T. nigerrimum*, while excluding taxa associated with stable, shaded microhabitats. Moderate disturbance can increase habitat heterogeneity at small spatial scales, but sustained or intensive disturbance tends to homogenise communities and reduce functional diversity (McKinney, 2008; Piano et al., 2020; Barton et al., 2024). These results highlight the dual role of disturbance as both a driver of local diversity and a filter excluding sensitive species, depending on intensity and frequency.

### 4.4. Species-specific effects of the invasive ant Linepithema humile

The invasive Argentine ant (*Linepithema humile*) was recorded in only three of the twelve studied green spaces, and its dominance varied markedly among them, reaching very high trap occupancy in the Botanical Garden (89.2%), moderate levels in CM-Asunción (25.4%), and relatively low values in Jardines de Renfe (15%). This restricted occurrence and variability in its presence suggests that *L. humile* success within the urban matrix depends strongly on local habitat conditions rather than urbanisation intensity alone (Holway et al., 2002; Abril, Oliveiras, et al., 2010; Abril et al., 2026).

Despite its restricted distribution, *L. humile* exerted strong, species-specific effects on native ant assemblages. Its presence was associated with significant declines in several dominant and widespread taxa (*L. grandis*, *P. pallidula*, *T. nigerrimum*, *T. semilaeve*, and *H. eduardi*; Table 2, Fig. 5), consistent with competitive exclusion driven by numerical dominance and interference competition (Carpintero et al., 2007; Madrzyk et al., 2026). In contrast, *Myrmica aloba* showed a strong positive association with *L. humile*, potentially reflecting indirect facilitation or shared tolerance to microclimatic conditions rather than direct ecological affinity (Lessard et al., 2009).

No significant relationships were detected for several other species (*C. mauritanica*, *P. pygmaea*, *A. gibbosa*, *A. senilis*, and *Solenopsis* spp.), indicating that some taxa may persist in invaded sites by exploiting less contested microhabitats or resources (Arnan et al., 2015; Blight et al., 2017; Trigos-Peral & Reyes López, 2016). These results highlight that invasive impacts operate through mechanisms partly independent of vegetation structure, reinforcing the need to evaluate invasion effects at the species and microhabitat level.

From an urban planning perspective, the strong but localised effects of *L. humile* underscore the importance of habitat-based management strategies. Structurally heterogeneous green spaces with diverse ground-layer features may enhance resistance to invasion and mitigate competitive exclusion, whereas simplified, resource-rich habitats appear more susceptible to invasive dominance (Menke et al., 2011; Suarez et al., 2005; Yamaguchi, 2004).

### 4.5. Implications for landscape and urban planning

Taken together, our results demonstrate that urban ant diversity – likely representative of broader patterns across other animal groups – is maximised not by increasing green cover alone, but by deliberately designing and managing urban green spaces to promote fine-scale structural and microhabitat heterogeneity. Biodiversity patterns arose from the interaction between vegetation structure, organic ground layers, and coarse structural elements, emphasising that urban green spaces should be viewed as multi-layered ecological systems rather than uniform vegetated areas.

Trees, leaf litter, stones, woody debris, and moderate disturbance regimes collectively supported diverse ant assemblages by providing complementary thermal refugia, nesting substrates, and foraging opportunities. In particular, fine-scale features such as stones, woody debris, and patches of bare ground generated thermally and structurally diverse microsites that functioned as key ecological resources. Although frequently removed during routine maintenance, these elements represent low-cost and effective means of increasing habitat heterogeneity within urban landscapes.

From a planning perspective, these findings translate into clear design principles: prioritising intermediate tree densities over uniform canopy closure, retaining leaf litter and dead wood where safety permits, incorporating stones and semi-natural substrates, and reducing the dominance of intensively managed lawns. Embedding these elements into urban green infrastructure can enhance ecological functionality, promote native biodiversity, and strengthen the role of cities within regional conservation networks.

Beyond their ecological importance, ants are also widely recognised as effective bioindicators of environmental quality and habitat condition in both natural and urban ecosystems. Their sensitivity to microclimatic variation, habitat disturbance, vegetation structure, soil conditions, and management intensity allows ant assemblages to respond rapidly to environmental change, often reflecting broader biodiversity patterns across other arthropod groups (Andersen, 1997; Underwood & Fisher, 2006). In urban landscapes, changes in ant community composition have been linked to habitat fragmentation, urban heat, vegetation simplification, and biological invasions (Menke et al., 2011; Carpintero & Reyes-López, 2014; Trigos-Peral et al., 2020; Lokatis & Jeschke, 2022; Trigos-Peral, Maák, et al., 2024; Trigos-Peral, Witek, et al., 2024; Trigos-Peral & Reyes-López, 2025; Penick & Edenborough, 2026; Czarnecka et al., 2026). The strong species-specific and microhabitat-dependent responses observed in our study further reinforce the value of ants as practical indicators of the ecological functionality of urban green spaces. Consequently, ant assemblages may provide a useful and cost-effective tool for assessing the success of biodiversity-oriented urban planning and management strategies aimed at increasing habitat heterogeneity and ecological resilience.

Overall, our results indicate that urban ecological design should move beyond vegetation quantity alone and instead promote heterogeneous microhabitat mosaics that support resilient arthropod assemblages while also reinforcing the use of ants as reliable tools for landscape evaluation and design, given their practical value as indicators of the ecological quality and functionality of urban green infrastructure.

## Statements and declarations

### Data Availability

Datasets and R codes used in the manuscript are available in figshare repository http://10.6084/m9.figshare.32326827. Datasets and R codes will be published online once the manuscript is accepted.

### Ethics approval

NA

### Patient consent

NA

### Permission to reproduce material from other sources

NA

### Clinical trial registration

NA

### Declaration of generative AI and AI-assisted technologies in the manuscript preparation process

During the preparation of this work the authors used Copilot AI in order to improve the aesthetics of the designed graphical abstract and to correct the English grammar. After using this tool/service, the author(s) reviewed and edited the content as needed and take(s) full responsibility for the content of the published article.

---

**Supplementary Figure 1.**
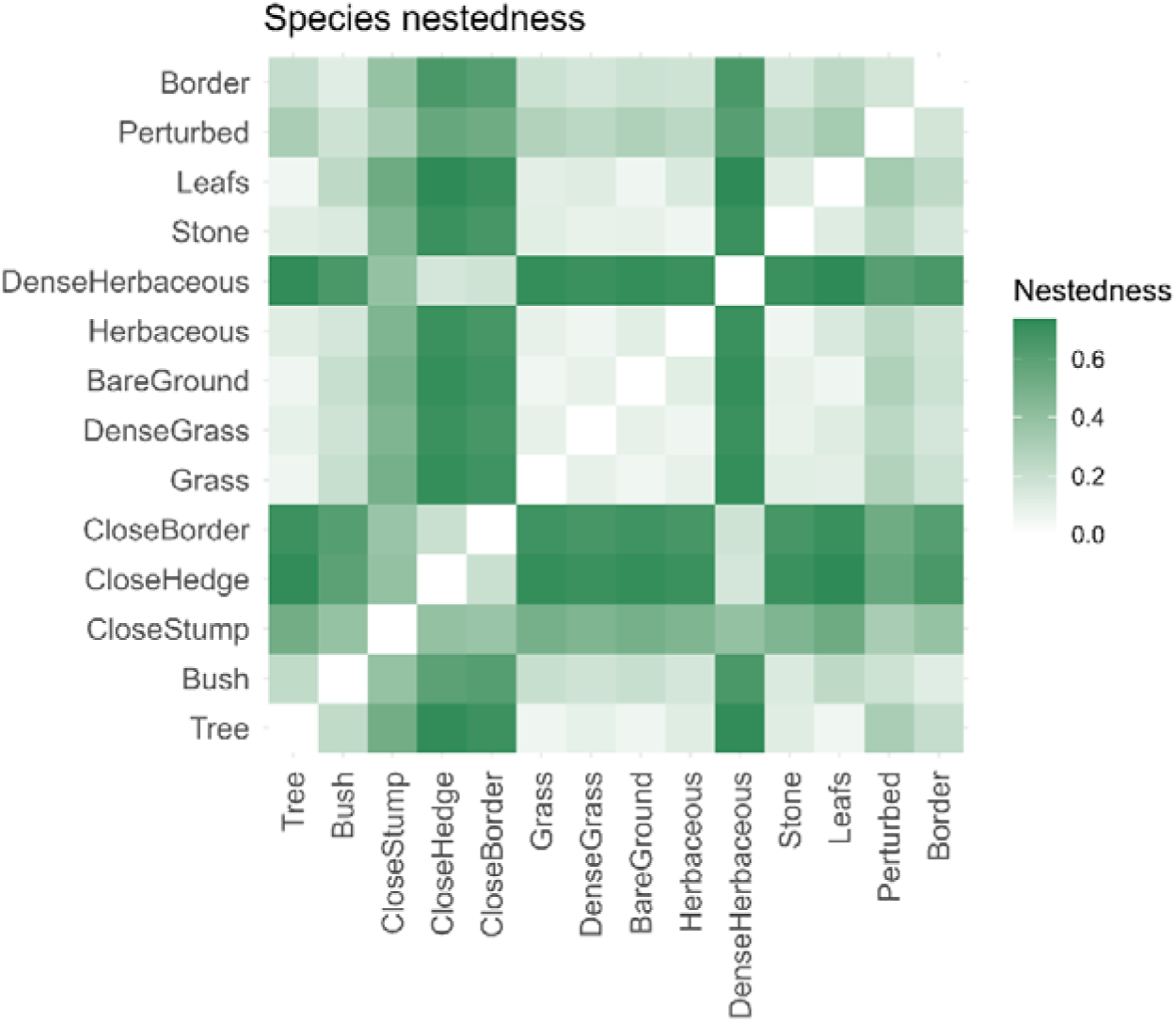
Heatmap showing Jaccard-based species nestedness among the different components of the microhabitat. Higher turnover is indicated by darker green squares.

**Supplementary Table 1.** Description of microhabitat components identified in the studied parks during the sampling period, based on information in situ and photographic analysis.

| Parameter | Description |
| --- | --- |
| Tree foot | The distance between the trap and the tree is less than or equal to twice the diameter of the top opening of the trap. |
| Under tree | Starting from a distance to the foot twice as large as the diameter of the upper opening of the trap, and it must also remain under the area of influence of the tree. |
| Bush foot | The distance between the trap and the bush is less than or equal to twice the diameter of the top opening of the trap. |
| Under bush | Starting from a distance to the foot twice the diameter of the upper opening of the trap, and it must also remain under the area of influence of the shrub. |
| Close to stump | The distance between the stump and the trap should be equal to or less than twice the diameter of the trap's top opening. |
| Close to hedge | The distance between the hedge and the trap should be equal to or less than twice the diameter of the top opening of the trap. |
| Close to border | The distance between the rim and the trap is equal to or less than twice the diameter of the top opening of the trap. |
| Grass | It is defined as when the grass is not dense and we can see part of the substrate. |
| Dense grass | In this case we assume that the substrate is not visible and there is no other type of vegetation. |
| Bare ground | This parameter is defined when there is no vegetation or leaf litter present. |
| Herbaceous | In this case we consider that there is only vegetation, or that it is mixed with grass, or that it is possible to observe the substrate. |
| Dense herbaceous | Dense vegetation is considered to exist when the soil is not visible. |
| Stone | This parameter is defined when some type of rock is observed in the area. |
| Leafs | This parameter is defined when leaf litter (either sparse or dense) is present. |
| Perturbed | When the area surrounding the trap shows noticeable changes with respect to the original situation, once the trap has been placed. |
| Border | Trap located on the grass-exposed soil edge of a tree pit or path. |

**Supplementary table 1.**
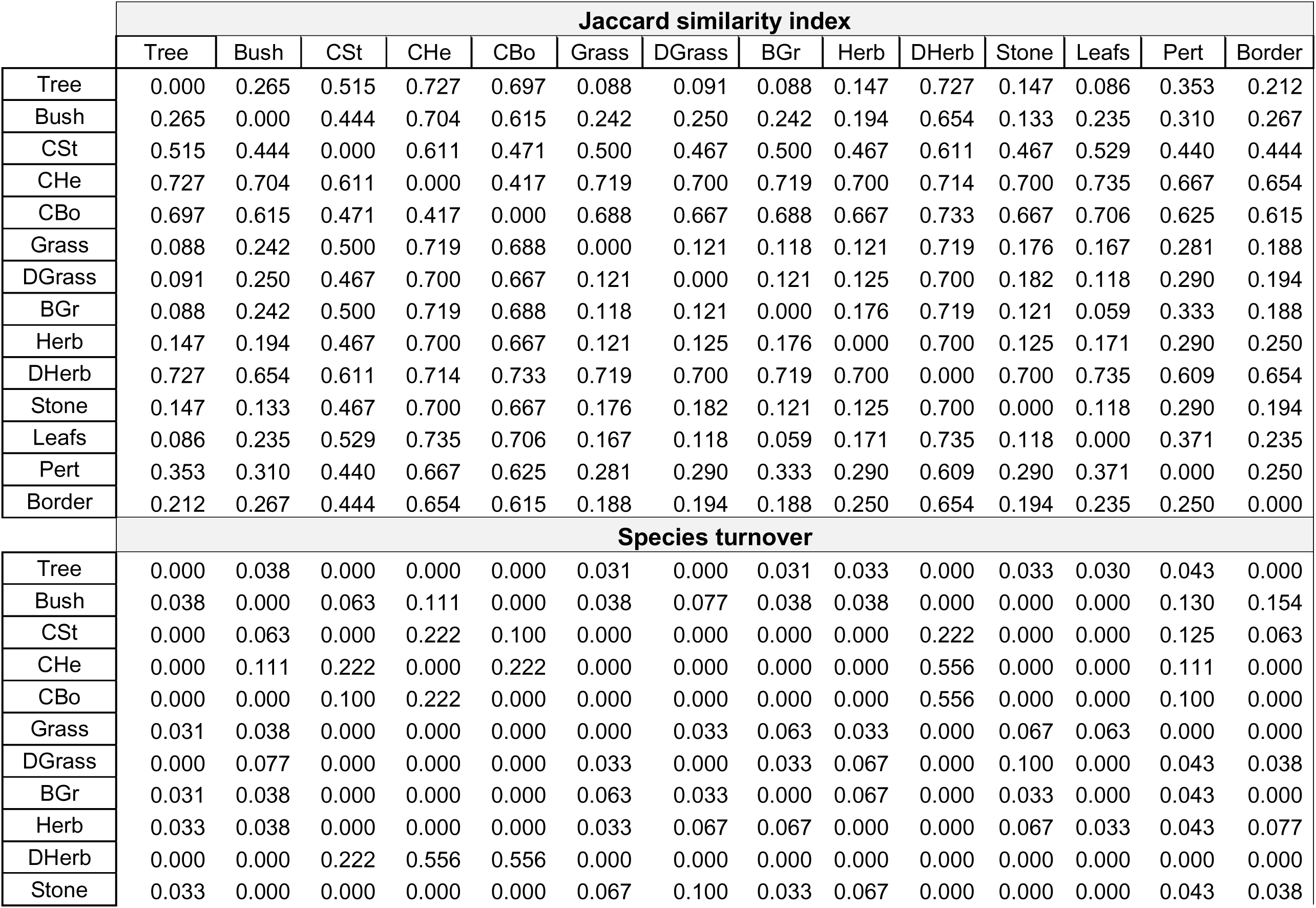
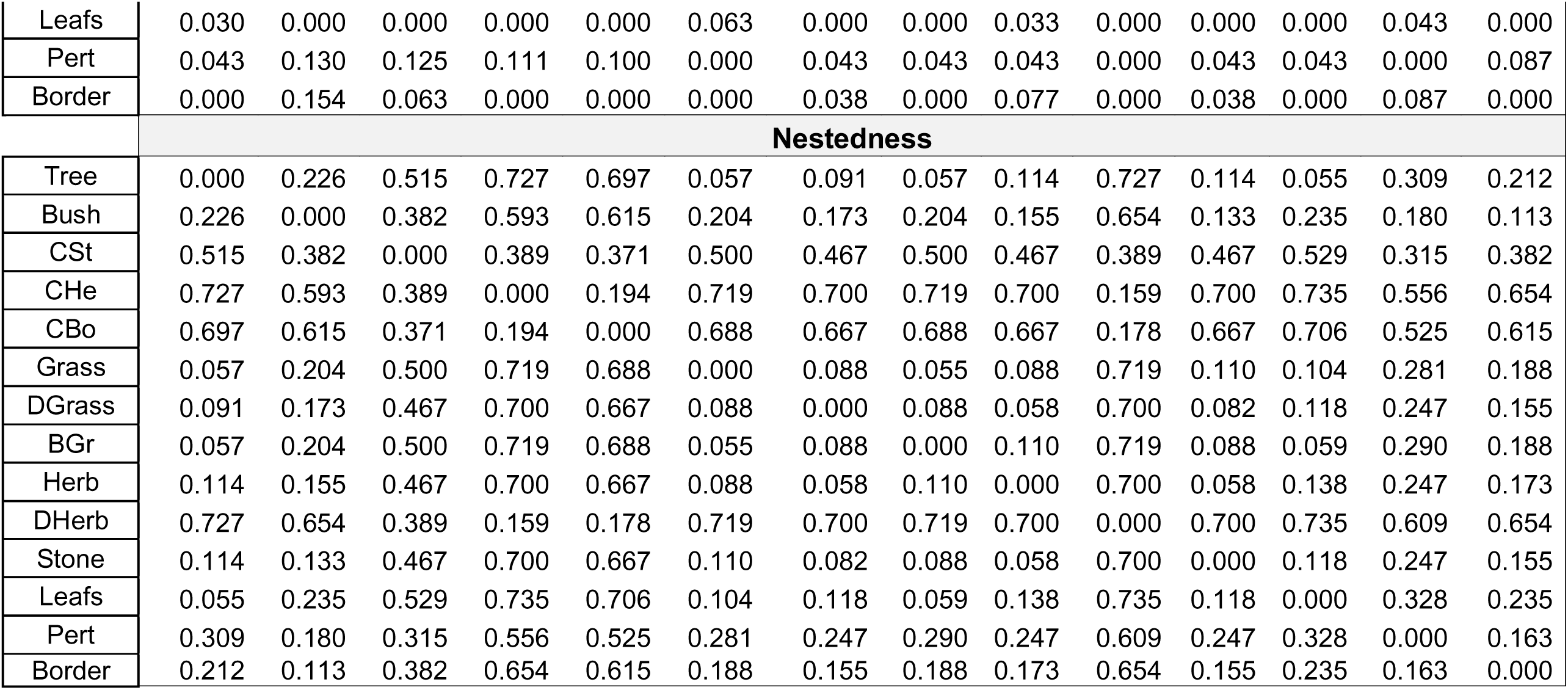
Jaccard similarity and its partitioning into species turnover and nestedness components among microhabitat elements. Values represent pairwise dissimilarities between microhabitat components. Effect sizes are interpreted as follows: values < 0.1 indicate low influence, values between 0.1 and 0.3 indicate moderate influence, and values > 0.3 indicate high influence on the studied index. Microhabitat components are: Tree, Bush, Close Stump (CSt), Close Hedge (CHe), Close Border (CBo), Grass, Dense Grass (DGrass), Bare Ground (BGr), Herbaceus (Herb), Dense Herbaceous (DHerb), Stone, Leaflitter (Leafs), Perturbed (Pert), Border.

Landscape and Urban planning
|  | Jaccard similarity index |  |  |  |  |  |  |  |  |  |  |  |  |  |
| --- | --- | --- | --- | --- | --- | --- | --- | --- | --- | --- | --- | --- | --- | --- |
|  | Tree | Bush | CSt | CHe | CBo | Grass | DGrass | BGr | Herb | DHerb | Stone | Leafs | Pert | Border |
| Tree | 0.000 | 0.265 | 0.515 | 0.727 | 0.697 | 0.088 | 0.091 | 0.088 | 0.147 | 0.727 | 0.147 | 0.086 | 0.353 | 0.212 |
| Bush | 0.265 | 0.000 | 0.444 | 0.704 | 0.615 | 0.242 | 0.250 | 0.242 | 0.194 | 0.654 | 0.133 | 0.235 | 0.310 | 0.267 |
| CSt | 0.515 | 0.444 | 0.000 | 0.611 | 0.471 | 0.500 | 0.467 | 0.500 | 0.467 | 0.611 | 0.467 | 0.529 | 0.440 | 0.444 |
| CHe | 0.727 | 0.704 | 0.611 | 0.000 | 0.417 | 0.719 | 0.700 | 0.719 | 0.700 | 0.714 | 0.700 | 0.735 | 0.667 | 0.654 |
| CBo | 0.697 | 0.615 | 0.471 | 0.417 | 0.000 | 0.688 | 0.667 | 0.688 | 0.667 | 0.733 | 0.667 | 0.706 | 0.625 | 0.615 |
| Grass | 0.088 | 0.242 | 0.500 | 0.719 | 0.688 | 0.000 | 0.121 | 0.118 | 0.121 | 0.719 | 0.176 | 0.167 | 0.281 | 0.188 |
| DGrass | 0.091 | 0.250 | 0.467 | 0.700 | 0.667 | 0.121 | 0.000 | 0.121 | 0.125 | 0.700 | 0.182 | 0.118 | 0.290 | 0.194 |
| BGr | 0.088 | 0.242 | 0.500 | 0.719 | 0.688 | 0.118 | 0.121 | 0.000 | 0.176 | 0.719 | 0.121 | 0.059 | 0.333 | 0.188 |
| Herb | 0.147 | 0.194 | 0.467 | 0.700 | 0.667 | 0.121 | 0.125 | 0.176 | 0.000 | 0.700 | 0.125 | 0.171 | 0.290 | 0.250 |
| DHerb | 0.727 | 0.654 | 0.611 | 0.714 | 0.733 | 0.719 | 0.700 | 0.719 | 0.700 | 0.000 | 0.700 | 0.735 | 0.609 | 0.654 |
| Stone | 0.147 | 0.133 | 0.467 | 0.700 | 0.667 | 0.176 | 0.182 | 0.121 | 0.125 | 0.700 | 0.000 | 0.118 | 0.290 | 0.194 |
| Leafs | 0.086 | 0.235 | 0.529 | 0.735 | 0.706 | 0.167 | 0.118 | 0.059 | 0.171 | 0.735 | 0.118 | 0.000 | 0.371 | 0.235 |
| Pert | 0.353 | 0.310 | 0.440 | 0.667 | 0.625 | 0.281 | 0.290 | 0.333 | 0.290 | 0.609 | 0.290 | 0.371 | 0.000 | 0.250 |
| Border | 0.212 | 0.267 | 0.444 | 0.654 | 0.615 | 0.188 | 0.194 | 0.188 | 0.250 | 0.654 | 0.194 | 0.235 | 0.250 | 0.000 |
|  | Species turnover |  |  |  |  |  |  |  |  |  |  |  |  |  |
| Tree | 0.000 | 0.038 | 0.000 | 0.000 | 0.000 | 0.031 | 0.000 | 0.031 | 0.033 | 0.000 | 0.033 | 0.030 | 0.043 | 0.000 |
| Bush | 0.038 | 0.000 | 0.063 | 0.111 | 0.000 | 0.038 | 0.077 | 0.038 | 0.038 | 0.000 | 0.000 | 0.000 | 0.130 | 0.154 |
| CSt | 0.000 | 0.063 | 0.000 | 0.222 | 0.100 | 0.000 | 0.000 | 0.000 | 0.000 | 0.222 | 0.000 | 0.000 | 0.125 | 0.063 |
| CHe | 0.000 | 0.111 | 0.222 | 0.000 | 0.222 | 0.000 | 0.000 | 0.000 | 0.000 | 0.556 | 0.000 | 0.000 | 0.111 | 0.000 |
| CBo | 0.000 | 0.000 | 0.100 | 0.222 | 0.000 | 0.000 | 0.000 | 0.000 | 0.000 | 0.556 | 0.000 | 0.000 | 0.100 | 0.000 |
| Grass | 0.031 | 0.038 | 0.000 | 0.000 | 0.000 | 0.000 | 0.033 | 0.063 | 0.033 | 0.000 | 0.067 | 0.063 | 0.000 | 0.000 |
| DGrass | 0.000 | 0.077 | 0.000 | 0.000 | 0.000 | 0.033 | 0.000 | 0.033 | 0.067 | 0.000 | 0.100 | 0.000 | 0.043 | 0.038 |
| BGr | 0.031 | 0.038 | 0.000 | 0.000 | 0.000 | 0.063 | 0.033 | 0.000 | 0.067 | 0.000 | 0.033 | 0.000 | 0.043 | 0.000 |
| Herb | 0.033 | 0.038 | 0.000 | 0.000 | 0.000 | 0.033 | 0.067 | 0.067 | 0.000 | 0.000 | 0.067 | 0.033 | 0.043 | 0.077 |
| DHerb | 0.000 | 0.000 | 0.222 | 0.556 | 0.556 | 0.000 | 0.000 | 0.000 | 0.000 | 0.000 | 0.000 | 0.000 | 0.000 | 0.000 |
| Stone | 0.033 | 0.000 | 0.000 | 0.000 | 0.000 | 0.067 | 0.100 | 0.033 | 0.067 | 0.000 | 0.000 | 0.000 | 0.043 | 0.038 |
| Landscape and Urban planning |  |  |  |  |  |  |  |  |  |  |  |  |  |  |
| Leafs | 0.030 | 0.000 | 0.000 | 0.000 | 0.000 | 0.063 | 0.000 | 0.000 | 0.033 | 0.000 | 0.000 | 0.000 | 0.043 | 0.000 |
| Pert | 0.043 | 0.130 | 0.125 | 0.111 | 0.100 | 0.000 | 0.043 | 0.043 | 0.043 | 0.000 | 0.043 | 0.043 | 0.000 | 0.087 |
| Border | 0.000 | 0.154 | 0.063 | 0.000 | 0.000 | 0.000 | 0.038 | 0.000 | 0.077 | 0.000 | 0.038 | 0.000 | 0.087 | 0.000 |
| Nestedness |  |  |  |  |  |  |  |  |  |  |  |  |  |  |
| Tree | 0.000 | 0.226 | 0.515 | 0.727 | 0.697 | 0.057 | 0.091 | 0.057 | 0.114 | 0.727 | 0.114 | 0.055 | 0.309 | 0.212 |
| Bush | 0.226 | 0.000 | 0.382 | 0.593 | 0.615 | 0.204 | 0.173 | 0.204 | 0.155 | 0.654 | 0.133 | 0.235 | 0.180 | 0.113 |
| CSt | 0.515 | 0.382 | 0.000 | 0.389 | 0.371 | 0.500 | 0.467 | 0.500 | 0.467 | 0.389 | 0.467 | 0.529 | 0.315 | 0.382 |
| CHe | 0.727 | 0.593 | 0.389 | 0.000 | 0.194 | 0.719 | 0.700 | 0.719 | 0.700 | 0.159 | 0.700 | 0.735 | 0.556 | 0.654 |
| CBo | 0.697 | 0.615 | 0.371 | 0.194 | 0.000 | 0.688 | 0.667 | 0.688 | 0.667 | 0.178 | 0.667 | 0.706 | 0.525 | 0.615 |
| Grass | 0.057 | 0.204 | 0.500 | 0.719 | 0.688 | 0.000 | 0.088 | 0.055 | 0.088 | 0.719 | 0.110 | 0.104 | 0.281 | 0.188 |
| DGrass | 0.091 | 0.173 | 0.467 | 0.700 | 0.667 | 0.088 | 0.000 | 0.088 | 0.058 | 0.700 | 0.082 | 0.118 | 0.247 | 0.155 |
| BGr | 0.057 | 0.204 | 0.500 | 0.719 | 0.688 | 0.055 | 0.088 | 0.000 | 0.110 | 0.719 | 0.088 | 0.059 | 0.290 | 0.188 |
| Herb | 0.114 | 0.155 | 0.467 | 0.700 | 0.667 | 0.088 | 0.058 | 0.110 | 0.000 | 0.700 | 0.058 | 0.138 | 0.247 | 0.173 |
| DHerb | 0.727 | 0.654 | 0.389 | 0.159 | 0.178 | 0.719 | 0.700 | 0.719 | 0.700 | 0.000 | 0.700 | 0.735 | 0.609 | 0.654 |
| Stone | 0.114 | 0.133 | 0.467 | 0.700 | 0.667 | 0.110 | 0.082 | 0.088 | 0.058 | 0.700 | 0.000 | 0.118 | 0.247 | 0.155 |
| Leafs | 0.055 | 0.235 | 0.529 | 0.735 | 0.706 | 0.104 | 0.118 | 0.059 | 0.138 | 0.735 | 0.118 | 0.000 | 0.328 | 0.235 |
| Pert | 0.309 | 0.180 | 0.315 | 0.556 | 0.525 | 0.281 | 0.247 | 0.290 | 0.247 | 0.609 | 0.247 | 0.328 | 0.000 | 0.163 |
| Border | 0.212 | 0.113 | 0.382 | 0.654 | 0.615 | 0.188 | 0.155 | 0.188 | 0.173 | 0.654 | 0.155 | 0.235 | 0.163 | 0.000 |

